# Hedgehog signaling can enhance glycolytic ATP production in the *Drosophila* wing disc

**DOI:** 10.1101/2021.09.11.459911

**Authors:** Ioannis Nellas, K. Venkatesan Iyer, Juan M. Iglesias-Artola, André Nadler, Natalie A. Dye, Suzanne Eaton

**Affiliations:** Max Planck Institute for Molecular Cell Biology and Genetics, Dresden, Germany; Department of Mechanical Engineering, Indian Institute of Science, Bangalore, 560012, India; Mildred Scheel Nachwuchszentrum (MSNZ) P2, Medical Faculty, Technische Universität Dresden, Germany; Excellence Cluster, Physics of Life, Technische Universität Dresden, Germany

**Keywords:** Hedgehog, glycolysis, ATP, *Drosophila*, metabolism

## Abstract

Energy production and utilization is critically important for animal development and growth. How it is regulated in space and time during tissue growth remains largely unclear. Toward this end, we used a FRET-based adenosine triphosphate (ATP) sensor to dynamically monitor ATP levels across a growing tissue, using the *Drosophila* wing disc. We discovered that steady-state levels of ATP are spatially uniform across the wing pouch. Pharmacologically inhibiting oxidative phosphorylation, however, reveals spatial heterogeneities in metabolic behavior, whereby signaling centers at compartment boundaries produce more ATP from glycolysis than the rest of the tissue. Genetic perturbations indicate that the conserved Hedgehog (Hh) signaling pathway can enhance ATP production by glycolysis. Collectively, our work reveals a positive feedback loop between Hh signaling and energy metabolism, advancing our understanding of the connection between conserved developmental patterning genes and energy production during animal tissue development.

## INTRODUCTION

The regulation of energy production and utilization is critically important for the growth of living organisms. The energetic currency ATP is produced by the breakdown of glucose to pyruvate during glycolysis (yielding 2 ATPs), followed by the oxidation of pyruvate through the TCA cycle and oxidative phosphorylation (OxPhos) under normoxic conditions (∼ 36 ATPs). In the absence of sufficient oxygen, pyruvate can be converted to lactate and secreted. Even under normoxic conditions, increased glucose uptake can result in the secretion of excess pyruvate as lactate, a phenomenon called “aerobic glycolysis”. This behavior is thought to allow intermediate metabolites from the glycolytic pathway and the TCA cycle to be channeled to other pathways to rapidly produce macromolecular precursors (nucleotides, amino acids, and lipids), which are also required for growth (Lunt and Vander Heiden, 2011).

Tissue growth during animal development also involves morphogens – secreted signals that pattern gene expression across a tissue in a concentration-dependent manner (Rogers and Schier, 2011). Hedgehog is one such morphogen that promotes the growth and patterning of many different tissues during development and also regulates many processes during adult homeostasis (Ingham and McMahon, 2001; Petrova and Joyner, 2014). One of the vertebrate Hh homologs, Sonic Hedgehog (SHH), has been shown to promote aerobic glycolysis in fat cells and in cerebellar granule neuron precursors (Teperino *et al*., 2012; Gershon *et al*., 2013; Di Magno *et al*., 2014). Thus, it is interesting to investigate how spatial gradients of Hh that form during tissue development may affect metabolism and energy production. Recently developed live, fluorescent biosensors have made it possible to address this question by enabling the monitoring of metabolites at high spatial-temporal resolution in living cells and tissues (Tsuyama *et al*., 2013; Bulusu *et al*., 2017; Greenwald, Mehta and Zhang, 2018; Volkenhoff *et al*., 2018).

Here, we study the spatial dynamics of ATP levels in the *Drosophila* wing disc, a growing tissue that has been a powerful model system for studying principles of morphogen signaling and developmental tissue growth (Hariharan, 2015; Beira and Paro, 2016). The patterns of morphogen signaling are well characterized, easy to visualize in the flat “pouch” region, and able to be genetically perturbed with spatial and temporal control. We exploit these properties to study how Hh may influence energy production. Previous work indicates that OxPhos is a major provider of ATP in the wing disc (Spannl *et al*., 2020) and that little aerobic glycolysis is normally observed in the tissue (De La Cova *et al*., 2014; Wang *et al*., 2016; Bawa *et al*., 2020). Nonetheless, glycolytic genes are expressed during normal wing disc development (Dye *et al*., 2017; Spannl *et al*., 2020) and their genetic depletion results in a mild but consistent reduction of steady state ATP levels and undergrowth of the tissue (Spannl *et al*., 2020). Furthermore, loss of glycolytic gene expression affects the plasma membrane potential, causing reduced uptake of Hh-inhibitory lipids and thereby an upregulation of Hh signaling. Thus, glycolysis has a function in maintaining Hh signaling and tissue growth in the wing disc. Whether Hh in turn affects energy production in this tissue is not known.

Using a recently developed FRET-based ATP sensor (Tsuyama *et al*., 2013), we dynamically monitored ATP levels across the wing disc tissue. We find that OxPhos inhibition reveals higher glycolytic ATP production in the Hh signaling domain and that genetic modification of the pathway can influence ATP production from glycolysis. Altogether, our work establishes positive feedback between morphogen signaling and metabolic activity during developmental tissue growth.

## RESULTS and DISCUSSION

### OxPhos inhibition reveals spatially heterogeneous metabolism in the wing pouch

We monitored ATP levels across the wing pouch in live explants using a ubiquitously expressed FRET-based ATP sensor, ubi-AT1.03NL (Fig 1A) (Tsuyama *et al*., 2013). Steady state levels of ATP are similar throughout the wing pouch and decline considerably upon OxPhos inhibition with antimycin A (Fig 1C, (Spannl *et al*., 2020)). After 2 hr of treatment with antimycin A, ATP levels fall close to the lower detection limit of the sensor (as determined by the use of the ATP-insensitive sensor ubi-AT1.03RK, Fig S1B & E-F), whereas ATP levels in control wing discs remain stable over this time period (Fig S1 C-D). Interestingly, a transient pattern emerges in the wing pouch during OxPhos inhibition (Fig 1C). ATP levels drop slightly slower in two perpendicular stripe regions, corresponding to the Dorsal-Ventral (DV) boundary and an area just anterior to the Anterior-Posterior (AP) boundary (Fig S1I-J). These areas are known growth “organizer” regions, with high signaling activity in the Wingless/Notch and Hedgehog pathways, respectively (Fig 1B).

**Figure 1:**
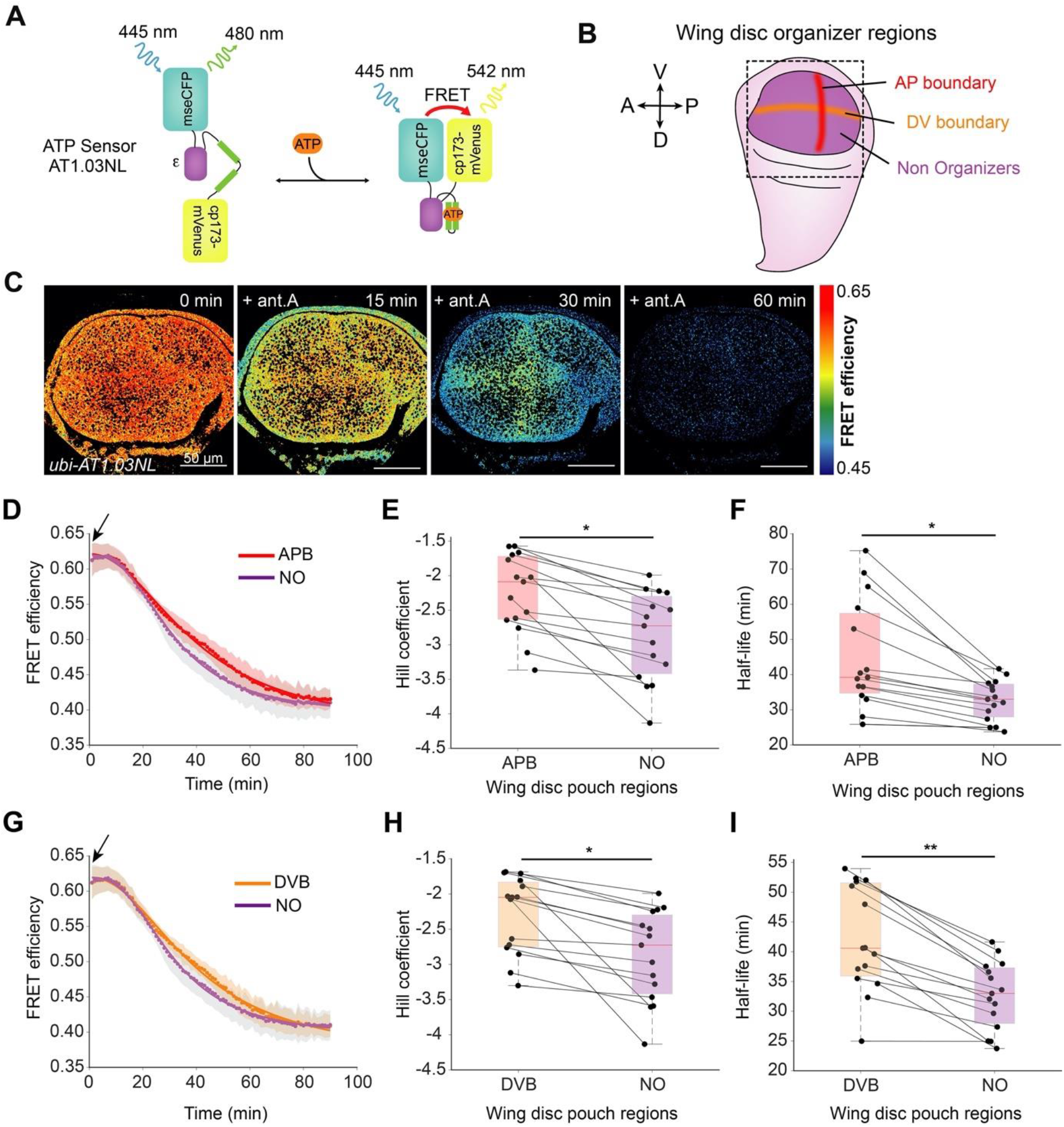
OxPhos inhibition depletes ATP levels slower at the organizers than elsewhere in the wing pouch. (A) Schematic representation of ATP FRET sensor design, adapted from (Tsuyama *et al*., 2013). Binding of ATP leads to a conformational change that brings the two fluorophores (donor and acceptor) in close proximity, resulting in FRET. (B) Schematic representation of the wing disc highlighting organizer and non-organizer regions. (C) Timelapse montage of ATP sensor FRET efficiency across the wing pouch after 10 μM antimycin A (ant.A) addition. (D, G) Mean FRET efficiency measured over time in the (D) AP boundary (APB) or (G) DV boundary (DVB) and non-organizer regions (NO). Shaded regions indicate standard deviation (SD); solid lines illustrate a fit to the mean data. (E-F, H-I) Fit parameters of individual time traces for AP boundary and non-organizer regions (E-F) or DV boundary and NO regions (H-I). Black lines connect the corresponding regions of the same disc. * = p-value < 0.05, ** = p-value < 0.01 using a Kruskal-Wallis test (n=15). Black arrows indicate the addition of the drug, and small colored dots represent the averages at each timepoint.

To quantitatively compare the kinetics of ATP depletion in different regions, we locally measured FRET efficiency over time during OxPhos inhibition (Fig 1D, G, Fig S1G-J). To describe the kinetic data, we used a four-parametric logistic curve that fits the data well. This fitting allows us to estimate the initial and final state of the FRET efficiency, the time needed to reach half of the FRET efficiency (half-life, units of time) and the slope of the curve at the half-life (Hill coefficient, unitless). We use the half-life and Hill coefficient to compare the kinetics of ATP depletion in different tissue regions (see Fig S1H and methods). Since the FRET efficiency decreases from a high state to a lower state, the Hill coefficient is negative. Consequently, a lower Hill coefficient corresponds to a steeper transition from the initial to the final state. In addition, lower values of half-life correspond to faster kinetics of ATP depletion. We compare relative changes in these kinetic fit parameters across different regions of the tissue. We find larger Hill coefficient and half-life values for the AP and DV boundary regions than for the rest of the pouch (Fig 1E-F, H-I). Thus, the AP and DV organizer regions have significantly slower kinetics of ATP depletion upon OxPhos inhibition than the rest of the wing pouch.

### Organizer regions can enhance glycolytic ATP production

ATP levels in organizer regions may decline slower upon OxPhos inhibition either because these regions consume ATP slower or produce more ATP from glycolysis (which is unaffected by antimycin A). To distinguish between these possibilities, we looked at the effect of simultaneously inhibiting glycolysis and OxPhos. We found that combining the OxPhos inhibitor antimycin A with the glycolytic inhibitors 3-bromo-pyruvate (3BP) and 2-deoxy-D-glucose (2DG) significantly alters the pattern observed with antimycin A alone (Fig 2A): ATP levels decay in organizer and non-organizer regions with similar kinetics (Fig 2B-D). Half-life values are indistinguishable (Fig 2D), and the Hill coefficient is slightly lower in the organizer regions than elsewhere in the tissue (Fig 2C). The latter indicates a slightly faster depletion of ATP in organizers, suggesting a higher ATP consumption rate. Thus, we conclude that organizer regions obtain more ATP from glycolysis during OxPhos inhibition than the rest of the wing pouch.

**Figure 2:**
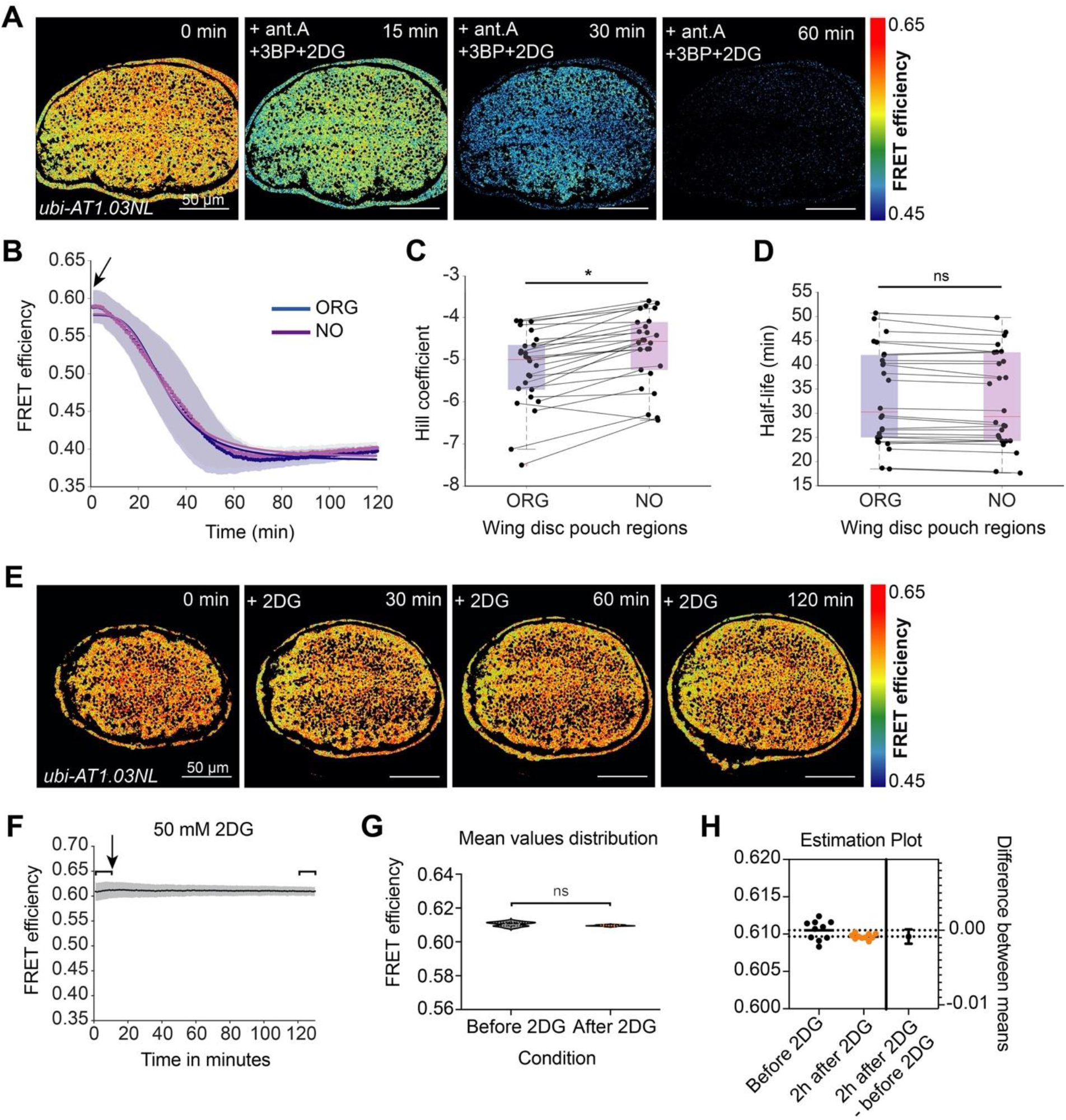
Glycolysis is required for spatial heterogeneity in ATP levels upon OxPhos inhibition. (A) Timelapse montage of ATP sensor FRET efficiency distribution after combined addition of 10 μM antimycin A (ant.A), 100 μM 3-Bromopyruvate (3BP), and 50 mM 2-deoxy-D-glucose (2DG). (B) Mean FRET efficiency of organizer (ORG) regions and non-organizer (NO) regions analyzed over time. Shaded regions indicate SD; blue and violet lines illustrate a fit of the mean data. (C, D) Fit parameters of individual time traces for organizer and non-organizer regions. Black lines connect the corresponding regions of the same disc. * = p-value < 0.05, ns = not significant p-value using a Kruskal-Wallis test (n=26). (E) Time lapse montage of ATP sensor FRET efficiency in the wing pouch after 2DG addition. (F)*Mean* FRET efficiency time trace of the entire wing pouch; gray shade indicates SD. Brackets include the mean FRET values before and 2h after 2DG addition (1-10 min and 121-130 min respectively) whose distribution was compared using (G) an unpaired t-test (ns = not significant p-value, n = 17) including (H) Welch’s correction and estimation plot (n=17 for each group). Black arrows indicate the addition of the drugs, and small colored dots represent the averages at each timepoint.

Note that the kinetics of ATP depletion throughout the wing pouch are faster upon inhibition of both OxPhos and glycolysis than upon inhibition of OxPhos alone, as indicated by a comparison of the Hill coefficients in Figs 1 and 2: -5.5 in non-organizers and -5.9 in organizers in all drugs (Fig 2C), compared to -2.7 in non-organizers and around -2.1 in both AP and DV boundaries in antimycin alone (Fig 1E, H). This finding indicates that upon OxPhos inhibition, glycolysis still generates ATP in the entire wing pouch but more so in the organizer regions. We were unable, however, to detect any difference in ATP levels with the FRET reporter after 2 hr of exposure to glycolytic inhibitors 3BP and 2DG alone or in combination, even when used at very high concentrations (Fig 2E-H, Fig S2D-I). Similarly, using a luciferase-based assay to measure ATP levels in whole wing discs, we did not find any significant effects of glycolytic inhibition (Fig S2A-B), even as OxPhos inhibition induced a rapid depletion of ATP (Fig S2C). We interpret these results to mean that most of the ATP in the wing disc is generated by OxPhos and that OxPhos can compensate for the 2 hr pharmacological inhibition of glycolysis. Nonetheless, the depletion of glycolytic enzymes by RNAi over several days results in a reduction of steady state ATP levels (Spannl *et al*., 2020). It remains possible that the FRET-ATP sensor is saturated and therefore cannot detect small drops in ATP upon glycolytic inhibition alone. We do not think this a likely scenario, however, given that most of our experiments start with an ATP FRET efficiency of only ∼0.6, but values of up to 0.7 can be observed in the same tissue under similar conditions, suggesting that the sensor can read higher concentrations of ATP.

### Hedgehog signaling enhances ATP production by glycolysis upon OxPhos inhibition

Our results suggest that the morphogen signaling that defines the organizer regions can influence energy production in the wing disc. It has been previously shown that Notch signaling, which is normally localized on either side of the DV boundary, can upregulate glycolysis (Slaninova *et al*., 2016), which is consistent with our data. The Hh pathway organizes growth and patterning along the AP axis, but its contribution to energy production has not been explored.

In the wing disc, Hh is expressed in the posterior compartment and travels to the anterior compartment, where it is received by the membrane protein, Patched (Ptc) (Basler and Struhl, 1994; Tabata and Kornberg, 1994). In the absence of Hh, Ptc represses the 7-pass signal transducer Smoothened (Denef *et al*., 2000). Hh binding to Ptc relieves this repression, allowing the Gli-family transcription factor Cubitus interruptus (Ci) to activate target gene expression.

To test whether Hh could affect energy production in the wing disc, we genetically activated and repressed the pathway in a temporally and spatially controlled manner using the Gal4/UAS system (Brand and Perrimon, 1993; Del Valle Rodríguez, Didiano and Desplan, 2012) and examined the effect on the kinetics of ATP depletion during pharmacological inhibition of glycolysis and OxPhos. We used *apterous-Gal4* combined with *tub-Gal80*^*ts*^ *(apGal*^*ts*^) to temporally perturb Hh signaling only in the dorsal compartment of the wing disc, leaving the ventral compartment as an internal control (Fig 3A). In the mock-treated *apGal*^*ts*^ genetic background alone (without a UAS construct), there is no difference in the kinetics of ATP depletion upon OxPhos inhibition between dorsal and ventral compartments (Fig S3).

**Figure 3.**
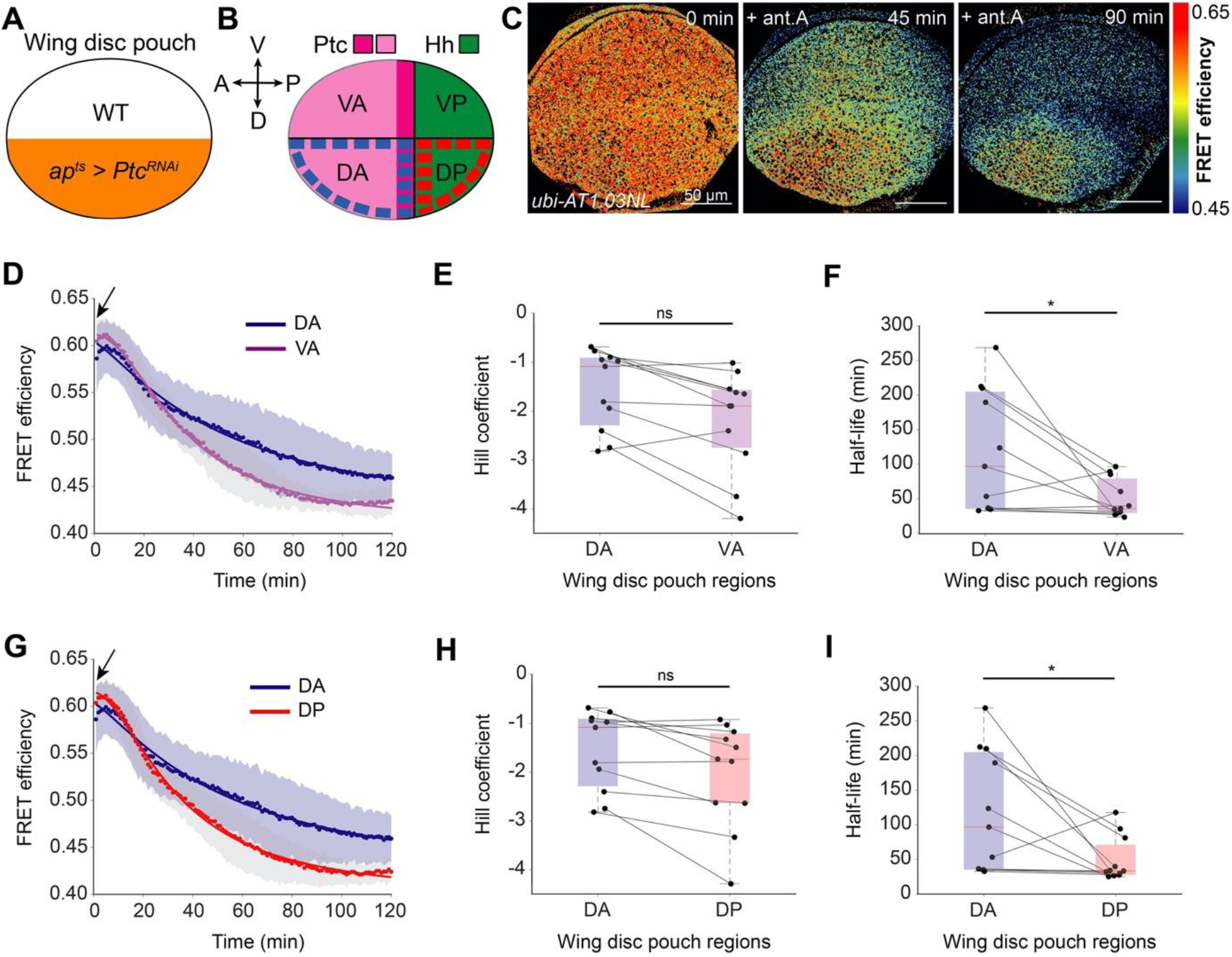
Upregulation of Hh activity enhances glycolytic ATP production upon OxPhos inhibition. (A) Schematic representation of Ptc knockdown in the dorsal compartment. (B) Schematic showing the expression pattern of Ptc and Hh in the wing disc. *Ptc* is expressed in the anterior compartment (light magenta) and its highest expression is at a narrow stripe right next to the AP boundary (dark magenta). (C) Timelapse montage of ATP sensor FRET efficiency in the wing pouch after 10 μM antimycin A (ant.A) addition in *apGal*^*ts*^>*Ptc*^*RNAi*^ wing discs. (D, G) Mean FRET efficiency analyzed over time in the (D) dorsal anterior (DA) and ventral anterior (VA) sub-compartments or the (F) dorsal anterior (DA) and dorsal posterior (DP) sub-compartments of *apGal*^*ts*^>*Ptc*^*RNAi*^ wing discs. Shaded regions indicate SD; solid lines illustrate a fit of the mean data. (E-F, H-I) Fit parameters of individual time traces for DA and VA compartments (E-F) or DA and DP compartments (H-I). Black lines connect the corresponding regions of the same disc. * = p-value < 0.05, ns = not significant p-value using a Kruskal-Wallis test (n=11). Black arrows indicate the addition of the drug and small colored dots represent the average values in each time point.

We used *apGal*^*ts*^ to overactivate the Hh pathway dorsally by inducing RNAi against *Ptc*, the Hh receptor that represses the pathway in the absence of ligand. *Ptc* is normally expressed in the anterior compartment, with a peak near the AP boundary (Fig 3B, S1J, S5A). Dorsal downregulation of Ptc does not affect the spatial pattern of steady state ATP levels in the wing disc but strikingly alters the pattern of metabolic activity in response to antimycin A (Fig 3C-I, Fig S4A-F). We observe a significantly longer half-life value for ATP depletion in the anterior region of the dorsal compartment (DA in Fig 3F, half-life of 96.7 min) compared to the anterior region of the ventral compartment (VA in Fig 3F, half-life of 33 min), which is not affected by *apGal*^*ts*^. The half-life for the DA region is also significantly longer than that of the posterior region of the dorsal compartment (DP, half-life of 33 min, Fig 3I). This difference can be explained by the fact that Ptc is normally only expressed in the anterior compartment. To confirm that this effect is cause by more ATP being produced by glycolysis, we again used the combination of OxPhos and glycolytic inhibitors (antimycin A, 3BP, and 2DG). Inhibiting both metabolic pathways eliminated the spatial differences in kinetics of ATP decline (Fig S4G-K). Thus, OxPhos inhibition reveals that overactivation of Hh signaling can enhance glycolytic ATP production.

Loss of Patched will increase the expression of another important growth regulator and Hh target gene, Decapentaplegic (Dpp). Dpp is produced in the anterior compartment in response to Hh signaling, but it is secreted and migrates bidirectionally through the tissue to promote growth and proliferation on both the anterior and posterior sides (Affolter and Basler, 2007; Restrepo, Zartman and Basler, 2014). Interestingly, in *apGal*^*ts*^>*Ptc*^*RNAi*^ discs, we found increased proliferation throughout the dorsal compartment, with no statistical difference between the anterior and posterior sides (Fig S5), consistent with an upregulation of Dpp. It is unlikely, however, that Dpp mediates the effect of Hh overactivation on glycolytic ATP production, as we only see an effect of *Ptc*^*RNAi*^ in the Hh-receiving anterior compartment, where Ptc is normally expressed (Fig 3B-I, Fig S4D-F).

To further test whether Hh pathway activity can regulate glycolysis in wing discs, we reduced Hh pathway activity by overexpressing a dominant negative form of the downstream Hh-responsive transcription factor, Cubitus interruptus (Ci^DN^, also named Ci^Cell^ in (Méthot and Basler, 1999)) (Fig 4). We used *apGal*^*ts*^ to induce Ci^DN^ in the dorsal compartment of the wing disc. This perturbation has the opposite effect of *Ptc*^*RNAi*^: upon loss of Hh activity, ATP levels decline faster during OxPhos inhibition (Fig 4C-E). Both the Hill coefficient and half-life values are significantly lower in the dorsal compartment than in the ventral (Hill coefficient = -4.5 for dorsal vs -1.8 for ventral; half-life = 32 min for dorsal, 42 min for ventral). This result suggests that down-regulation of Hh pathway activity using a dominant negative version of its downstream transcription factor reduces ATP production from glycolysis. Note that this perturbation, unlike *Ptc*^*RNAi*^, affects both the anterior and posterior sides of the dorsal compartment (Fig 4B). Although Ci is only expressed in the anterior compartment in wild type discs, here we are forcing the expression of a dominant negative construct in the entire dorsal domain, and therefore it is not unexpected to see an effect on both sides of the AP boundary.

**Figure 4.**
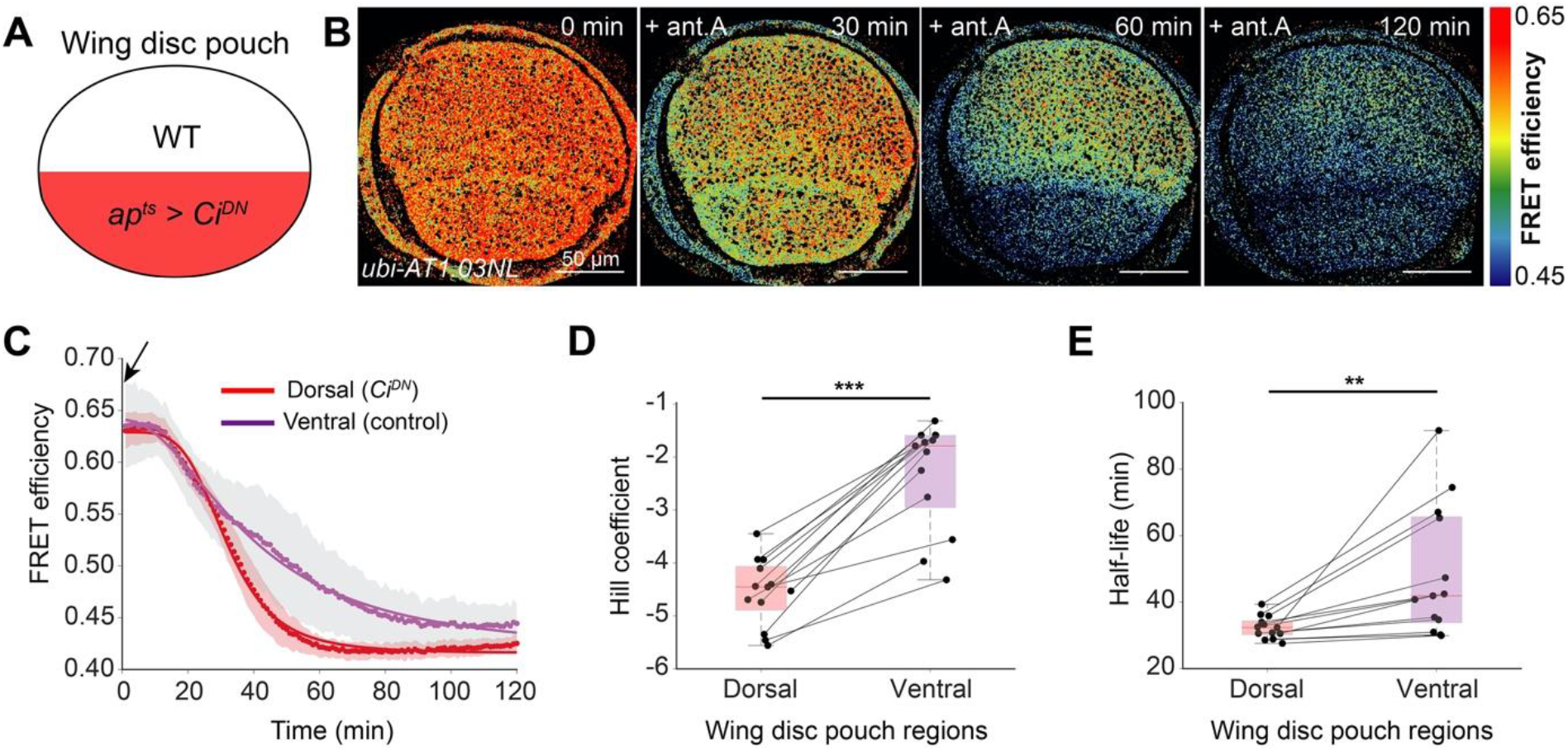
Downregulation of Hedgehog activity reduces glycolytic ATP production upon OxPhos inhibition. (A) Schematic showing the dorsal expression of the dominant negative Ci allele (Ci^DN^). (B) Timelapse montage of ATP sensor FRET efficiency distribution after 10 μM antimycin A addition in *apGal*^*ts*^ > *Ci*^*DN*^ wings. (C) Mean FRET efficiency measured over time in the dorsal and ventral compartments in *apGal*^*ts*^ > *Ci*^*DN*^ wings. Shaded regions indicate SD; solid lines illustrate a fit of the mean data. Black arrows indicate the addition of the drug, and small colored dots represent the averages at each timepoint. (D, E) Fit parameters of individual time traces for dorsal and ventral compartments. Black lines connect the corresponding regions of the same disc. ** = p-value < 0.01, *** = p-value < 0.001 using a Kruskal-Wallis test (n=13).

Taken together, our results from reciprocal genetic perturbations strongly suggest that Hh can positively affect glycolytic ATP production, at least upon inhibition of OxPhos. Given that glycolysis is required for normal levels of Hh signaling (Spannl *et al*., 2020), these new data indicate that there exists a positive feedback loop between Hh and glycolysis that could be part of a homeostatic mechanism coupling energy with developmental patterning and growth. Deciphering the underlying mechanism by which Hh promotes glycolysis in this model tissue is an important open question.

In summary, we show here that OxPhos inhibition reveals spatial heterogeneity in energy production in the wing pouch and that genetic perturbation experiments support a role for Hh in promoting ATP production by glycolysis. This work provides a foundation for new studies exploring how morphogen signaling and energy production interact to promote growth during tissue development.

## MATERIALS AND METHODS

### Fly stocks, husbandry and genetics

The following fly stocks were used: Wild-type Oregon-R (BDSC #5), *ap-Gal4* (BDSC #3041), *tub-Gal80*^*ts*^ (BDSC #7017 or 1019), *UAS-Ptc*^*RNAi*^ (BDSC) and *UAS-Ci*^*DN*^ (Ci^Cell^)(Méthot and Basler, 1999). All flies and larvae were raised on a standard food containing cornmeal, agar, malt, sugar beet syrup, brewery yeast, propionic acid and soy flour under a 12 hr light/dark cycle. All the knockdown and over-expression experiments for immunostainings and FRET analysis were performed with *apGal4-Gal80*^*ts*^ (used here as *apGal*^*ts*^). 20-30 female *apGal*^*ts*^ flies were crossed to male *UAS-Ptc*^*RNAi*^ *or UAS-Ci*^*DN*^ flies in a 3:1 ratio in a normal food vial. Flies were allowed to lay eggs in this vial in a 20 °C incubator or water bath and then were transferred to a new vial every day. Larval growth took place at 20 °C for one week, and then larvae were transferred to 30 °C for 24 hr. Then, upcrawling larvae were selected and both sexes were dissected. In all cases, the necessary controls (outcrosses with wild-type Oregon-R flies) were handled in the same way.

### Generation of transgenic lines

The FRET sensor ubi-AT1.03NL was generated as described in (Spannl *et al*., 2020). Additionally, *apGal*^*ts*^ was introduced into the background of the ubi-AT1.03NL flies and was used for the spatial-temporal overexpression of *Ptc*^*RNAi*^ and *Ci*^*DN*^.

### Imaging FRET-based ATP sensor in wing explants

Wing discs from upcrawling third-instar larvae were dissected in full medium (Grace’s medium (Sigma G9771) supplemented with 5% FBS (ThermoFischer/Invitrogen 10270098) and 20 nM of 20-hydroxyecdysone (Sigma H5142)(Dye *et al*., 2017) within 10 min and mounted as in (Spannl *et al*., 2020): basal side up on glass-bottom dishes (MatTek Corporation, #P35G-1.0-20 C) with a double-sided tape spacer and immobilized with a Whatman™ Cyclopore™ track-etched polycarbonate membrane filter (GE Healthcare Life Sciences, #7062-2513). Then, 1 ml of full medium was added and samples were transferred to the microscope. All experiments were performed at 25 °C, including the ones with *Ptc*^*RNAi*^, as we assume that new expression of Ptc will require longer than the 2 hr experiment.

For the experiments with metabolic drugs, 1 ml of full medium with 2X concentration of antimycin A (Sigma-Aldrich #A8674), 3-bromopyruvate (Sigma Aldrich #16490) or 2-deoxy-D-glucose (CARLROTH #CN96.3) was further added to the dish on the microscope using a hole on the lid. Drugs were added (shown with a black arrow in figures) either immediately after acquiring the first time point (time point 0 min) or after 10 min. Final concentrations are described in the figure legends.

Images of the wing pouch were acquired on an Olympus IX81 microscope equipped with CSU-W1 spinning disk (Yokogawa), Andor iXon Ultra 888, Monochrome EMCCD camera, Prior PRO SCAN III, Prior NanoScanZ and an incubation chamber to ensure stable temperature (25 °C). For all experiments a 60× silicone oil immersion objective lens was used (UPLSAPO60xS2, NA = 1.3). Wing discs were excited with a 445 nm laser twice in a sequential manner. Emission of mse-CFP was collected upon first excitation using an HQ 480/40 bandpass filter, and emission of cpVenus-FRET was collected using an HQ 542/27 filter. The bleedthrough of mse-CFP into the HQ 542/27 filter was estimated by exciting wing discs expressing ubi-Gal4-driven CFP-tagged human cytoplasmic β-actin (BDSC #7064) and acquiring images through an HQ 480/40 filter (I_D_) and its bleedthrough in an HQ 542/27 filter (I_bth_). The fraction of FRET intensity contributed by bleed-through is given by:

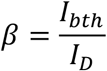

### Analysis of FRET-based ATP sensor imaging

FRET data were analyzed as previously described (Spannl *et al*., 2020). Briefly, a custom-written MATLAB (MathWorks) script was used to estimate the FRET efficiency from the fluorescence images after smoothening both donor and FRET images using a 5 × 5 averaging kernel (Spannl *et al*., 2020). Donor (*I*_*D*_) and FRET (*I*_*F*_) images were background subtracted, and the FRET intensity was corrected for bleedthrough as:

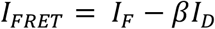

Finally, the FRET efficiency (η) was calculated as:

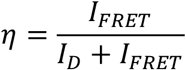

To separately quantify ATP dynamics in regions of the wing disc (Fig 1-3), the MATLAB script was modified by the Image Analysis Clinic of MPI-CBG to calculate FRET efficiency in user-defined circular ROIs of 20 μm diameter (Fig S1H). Using the maximum intensity projection of the cpVenus channel (not the FRET), it is possible to discern AP and DV boundary regions by eye (Fig S1G). The AP boundary appears as a stripe of slightly lower intensity, and the DV boundary lies in the middle of two strips of higher intensity. Using the MATLAB script, we define circular 9 ROIs (Fig S1H). The organizer region of the AP boundary was defined as the average of three circular ROIs in the AP compartment boundary, whereas the organizer region of the DV boundary was defined as the average of three circular ROIs in the DV compartment boundary. AP and DV boundaries shared the central ROI. Consequently, the average FRET efficiency of both organizer regions was estimated from these five circular ROIs described in the AP and DV compartment boundaries. Non-organizer regions were defined as the average of four circular ROIs laying outside of the AP and DV compartment boundaries (Fig S1G-H). To quantify FRET efficiency over the entire dorsal or ventral compartment of each disc (Fig 4, S3, S4A and S4G-K), a freehand tool was used to select these regions and calculate their average FRET efficiency.

### Empirical fit of ATP decline

Data from the FRET efficiency decline over time upon addition of metabolic drugs were fit using MATLAB (R2021a, The MathWorks Inc.) with the following equation:

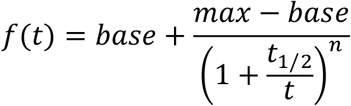

where *t*_1/2_ and n are the two fit parameters corresponding to the half-life (loss of 50% of FRET efficiency) and the Hill coefficient, respectively. Base and max were incorporated to account for the fact that our samples have variable starting and ending values of FRET. The base was set to be 0.2827, as this is the lowest reliable recorded FRET efficiency value derived from the z-stack of a morphologically healthy, unperturbed wing disc expressing ubiquitously the AT1.03RK sensor, as explained in Fig S1F. Max is also a fit parameter that never exceeded 0.7, which is the highest reliable mean FRET efficiency value that we have recorded, derived from morphologically healthy, unperturbed wing discs expressing ubiquitously the AT1.03NL sensor. Graphs were generated from the same script.

### Measurement of bulk ATP levels using a luciferase-based assay

Bulk levels of ATP were measured using a luciferase-based biochemical assay (ATPlite Luminescence Assay System, PerkinElmer). Wing discs from third instar upcrawling larvae were dissected in culture medium, washed with PBS within seconds, suspended in 20 μl of PBS, and added to wells of white polysterene flat bottom 96-well assay plates (Costar® 3917) containing 80 μl of PBS (to make a final volume of 100 μl in PBS). Specifically for Fig S2A-B, experiments were done with paired discs from each larva (right and left). One disc was treated with either 3BP or antimycin A (Fig S2A) or 3BP and 2DG (Fig S2B) and the other was mock-treated, serving as a control. The blank control was 100 μl of PBS. Samples were lysed by the addition of 50 μl of mammalian cell lysis solution followed by shaking at 700 rpm for 10 min. Luciferase substrate solution was added in a volume of 50 μl, and the samples were shaken at 700 rpm for 5 min. After 10 min of incubation in the dark, luciferase activity was measured using a Perkin Elmer Envision plate reader. To estimate concentration, a standard curve of luciferase activity was generated using a serial dilution of a 10 mM ATP stock solution.

To calculate the average wing disc volume, dissected wing discs from upcrawling larvae expressing the AT1.03NL FRET sensor were mounted and placed on the Olympus IX81 microscope as described earlier (Imaging FRET-based ATP in wing explants). Wing discs were excited with a 445 nm laser and emission of cpVenus-FRET was collected using an HQ 542/27 filter. Serial acquisition of images every 0.5 μm from the most apical to the most basal disc part resulted in a z-stack including the entire wing disc. The Image Analysis Clinic provided a FIJI macro that processes the z-stacks based on the fluorescence intensity (emission collected using the HQ 542/27 filter) and calculates the wing disc volume. The volume of 14 wing discs was used to estimate the average wing disc volume. To estimate the intracellular ATP concentration in approximation, the amount of ATP measured from single-discs luciferase assays was divided by the average wing disc volume.

### Immunofluorescence

Wing discs from upcrawling third instar larvae were dissected in PBS, fixed in 4 % paraformaldehyde (PFA) for 20 min, and rinsed three times in PBS. Wing discs were then permeabilized with 0.05 % Triton X-100 in PBS (PBX) twice for 10 min, blocked for 45 min in PBX + 1 mg/ml BSA + 250 mM NaCl (BBX 250), and incubated overnight with the primary antibody in PBX + 1 mg/ml BSA (BBX) at 4 ºC. After washing twice for 20 min in BBX, wing discs were blocked for 45 min in the blocking solution (BBX + 4 % normal goat serum) and incubated for 2-3 hr with the secondary antibody in the blocking solution. Afterwards, the wing discs were rinsed two times and washed three times within 45 min in PBX and the same in PBS. Finally, wing discs were mounted in VectaShield® (Vector Labs, #H-1000). Secondary antibodies conjugated with Alexa Fluor® 488 and 555 were diluted 1:1000 and Alexa Fluor® 647 were diluted 1:500 (ThermoFisher Scientific). Primary antibodies used:

| Antigen | Host species | Dilution used | Origin | Genotype |
| --- | --- | --- | --- | --- |
| Ptc | mouse | 1:100 | DSHB, AB_528441 | *apGal<sup>ts</sup>>Ptc<sup>RNAi</sup>/WT* |
| Ci<sub>155</sub> | rat | 1:30 | DSHB, AB_2109711 | *apGal<sup>ts</sup>>Ptc<sup>RNAi</sup>/WT* |
| PH3 | rabbit | 1:500 | Cell signaling #9701S | *apGal<sup>ts</sup>>Ptc<sup>RNAi</sup>/WT* |

Images were acquired using a Zeiss LSM700 inverted confocal microscope equipped with a Zeiss Axio Observer.Z1, a motorized stage Maerzhauser Wetzler Gmbh EK 130×85 mot. Tango CZ EMV, a 25×/0.8 LCI Plan-Neofluor, W/Glyc/Oil objective (Zeiss) and 2 PMT. Both samples and controls were dissected, fixed, stained, and imaged in parallel so that the reagents and handling conditions were always the same. Fiji (Schindelin *et al*., 2012) was used for image processing, orienting, and segmenting. Z-stacks of PH3 were max-projected and segmented using the Weka segmentation plugin (Arganda-Carreras *et al*., 2017) as previously described (Dye *et al*., 2017). Statistical analyses and plots were made using GraphPad Prism 9.

To confirm the colocalization of the FRET pattern close to the AP boundary with the Ptc stripe (Fig S1I-J), FRET efficiency was calculated in wing discs while treated with 10 μM antimycin A for up to 60 min, as previously described (Imaging FRET-based ATP in wing disc explant). At the end of 60 min, the wing discs were fixed on site with 4% PFA, and the glass bottom plates with the samples were removed so that immunostainings for the detection of Ptc with Alexa Fluor® 647 could continue, as described above. At the end of the immunostainings, the samples were taken back to the spinning disc microscope. Wing discs were excited with a 638 nm laser, and emission of Alexa Fluor® 647 was collected with a HQ 685/40 filter. Afterwards, images from both FRET and immunofluorescence experiments were compared for the same wing discs (Fig S1I-J).

### Statistical analyses

Statistical analyses were performed using GraphPad Prism 9 or MATLAB (R2021a, The MathWorks Inc.). For statistical significance, Kruskal-Wallis, paired t-tests, Mann Whitney tests, one-way ANOVA with Bonferroni correction and unpaired t-tests with Welch’s correction were performed as listed in the figure legends for each experiment. The use of either parametric or non-parametric statistical analysis was determined by the normal or not normal distribution of data, respectively.

## ACKNOWLEDGEMENTS

This work was supported by funding from the Max Planck Society and the Deutsche Forschungsgemeinschaft (SFB/TRR83). We thank the facilities of the MPI-CBG, specifically the Light Microscopy Facility, Scientific Computing (in particular Gayathri Nadar & Noreen Walker from the Image Analysis Clinic), and the fly facility (Sven Ssykor and Cornelia Maas). We also thank Suhrid Ghosh, Romina Piscitello-Gómez, Jonathan Rodenfels, and Michele Solimena for critical comments on the manuscript prior to submission. We heartfully thank the members of the Eaton lab, in particular Ali Mahmoud for excellent lab management and support. Lastly, we dedicate this paper to our inspiring mentor and co-author, Professor Dr. Suzanne Eaton, who tragically passed away near the conclusion of this project.

## AUTHOR CONTRIBUTIONS

IN performed experiments and analyzed data. KVI established the FRET acquisition methods, and JMIA developed the algorithmic tools used by IN to fit and analyze the FRET data. SE conceived the study. IN, SE, NAD and AN designed the study and analyzed data. IN and NAD wrote the manuscript with discussions and feedback from all authors.

## CONFLICT OF INTEREST

The authors declare no conflicts of interest.

---

**Extended data S1.**
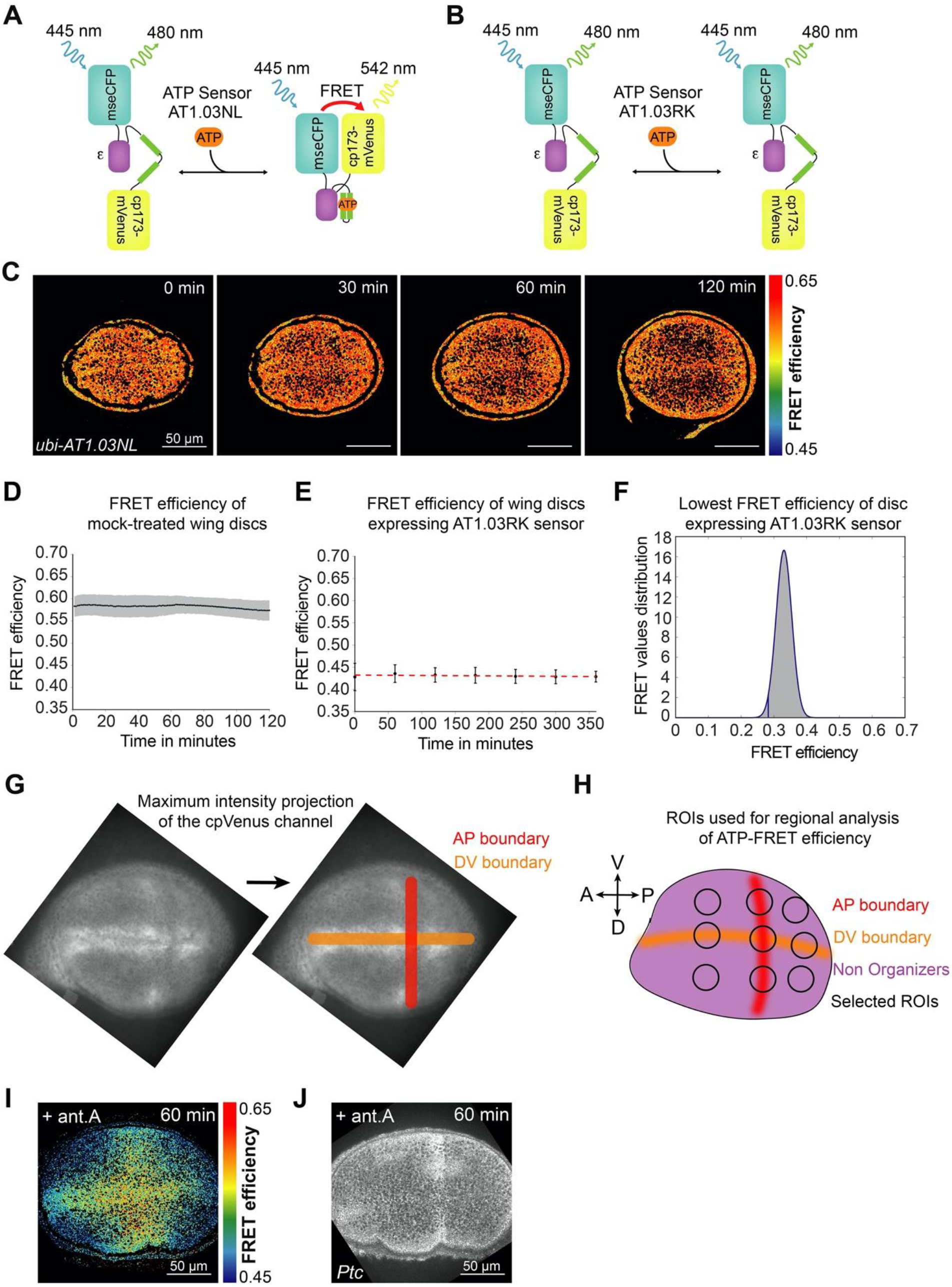
ATP levels in the wing pouch are spatially uniform and stable in culture without OxPhos inhibition. (A, B) Schematic representation of ATP-FRET sensor design (AT1.03NL) and its ATP-insensitive version (AT1.03RK). (C) Timelapse montage of ATP-FRET sensor efficiency in the wing pouch during culture for 2 hr. (D) Mean FRET efficiency in the entire pouch measured over time; gray shade indicates SD, and small black dots represent the averages at each time point (n=9). (E) Mean FRET efficiency in the entire pouch expressing AT1.03RK (red dashed line). The black error bars represent SD for each time point (n=7). (F) Histogram showing the distribution of FRET efficiency values across a 70-plane z-stack of a morphologically healthy, unperturbed wing disc ubiquitously expressing the AT1.03RK sensor: gray area corresponds to the FRET values, and the blue curved line shows a Gaussian fit. The lowest FRET efficiency value used for data fitting was 0.2827 (blue vertical line), defined as the mean FRET efficiency minus twice the standard deviation. (G) AP and DV boundary regions are discernible with a maximum intensity projection of the cpVenus channel. The AP boundary appears as a stripe of slightly lower intensity, and the DV boundary lies in the middle of two strips of higher intensity. (H) Schematic indicating the location of the ROIs that were used to calculate mean FRET efficiency in different wing pouch regions. The three ROIs in the AP boundary or in the DV boundary were averaged together to calculate the mean FRET efficiency of the AP boundary and DV boundary, respectively. The organizer region was measured as the average of all five of these ROIs (AP+DV boundaries). The non-organizer region corresponds to the remaining four ROIs outside of the AP+DV boundaries. (I) Spatial pattern of FRET efficiency after 60 min of 10 μM antimycin A (ant. A) exposure, and (J) Ptc expression in the same disc after fixation and immunofluorescence.

**Extended data S2.**
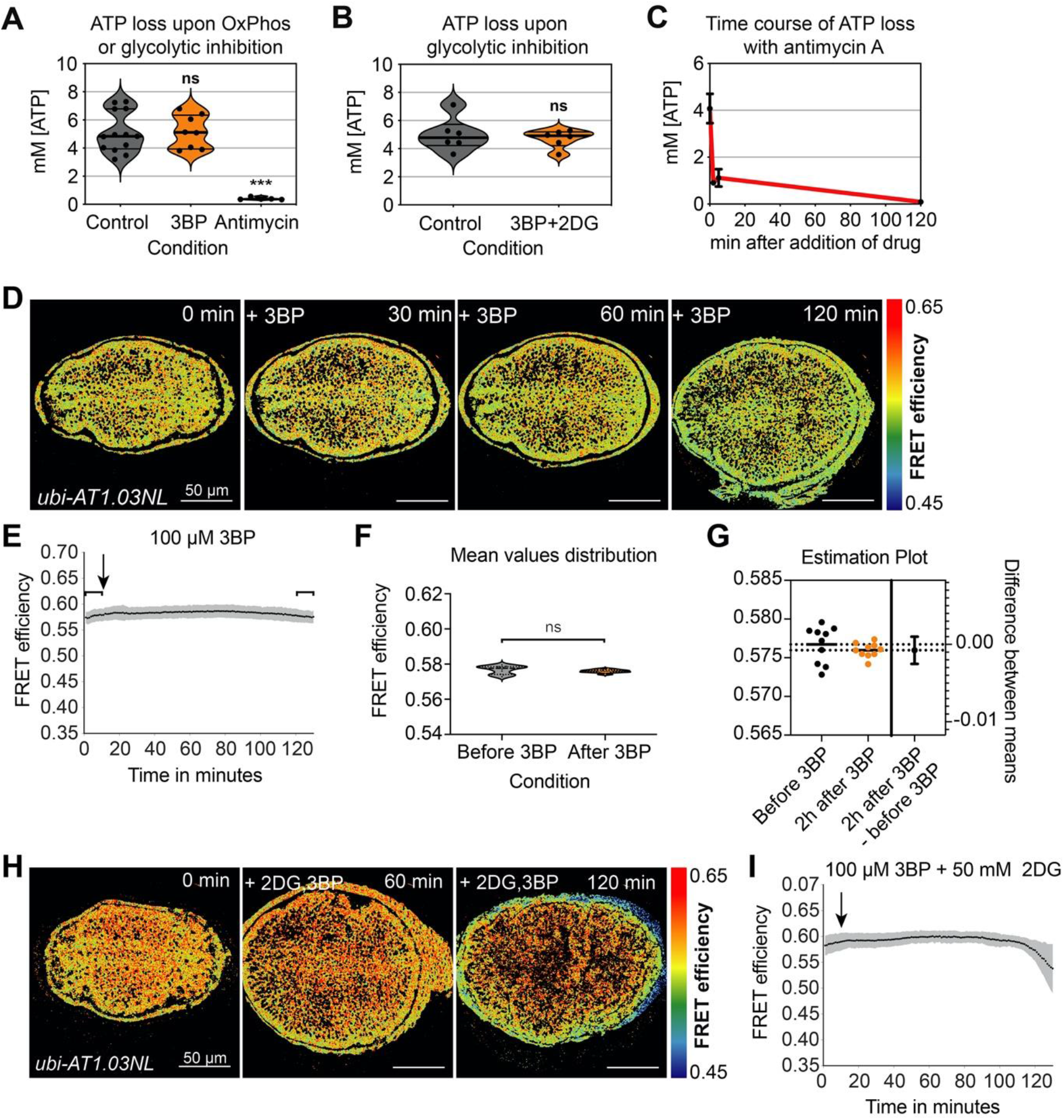
Glycolysis inhibitors alone do not significantly affect ATP levels. (A) ATP levels measured using a luminescence-based biochemical assay from single discs after a 2 hr treatment with either 50 μM 3BP (n=8) or 10 μM antimycin A (n=5) compared to untreated discs (n=13). *** = p-value < 0.001, ns = not significant using either Mann-Whitney test (control vs 3BP) or unpaired t-test (control vs antimycin A). (B) ATP levels of single discs after 1 hr of treatment with 50 μM 3BP + 50 mM 2DG (n=6) compared to untreated discs (n=6). ns = not significant using a paired t-test. (C) Decline of ATP levels upon addition of 100 μM antimycin A for varying lengths of time (n=10 for 0 min, n=3 for 2 min and 5 min and n=4 for 120 min). (D) Timelapse montage of ATP-FRET sensor efficiency after 3BP addition. (E) Mean FRET efficiency measured over time in the entire wing pouch upon addition of 3BP. Brackets include the mean FRET values before and 2 hr after 3BP addition (1-10 min and 121-130 min respectively) whose distribution was compared using (F) an unpaired t-test (ns = not significant p-value, n = 11) including (G) Welch’s correction and estimation plot (n = 11 for each group). (H) Timelapse montage of ATP-FRET sensor efficiency after 3BP+2DG addition. (I) Mean FRET efficiency measured over time in the entire wing pouch upon addition of 2DG+3BP; (E, I) gray shade indicates SD (n = 11 in (E) and n = 9 in (I)). Black arrows indicate the addition of the drugs, and small black dots represent the averages at each timepoint.

**Extended data S3.**
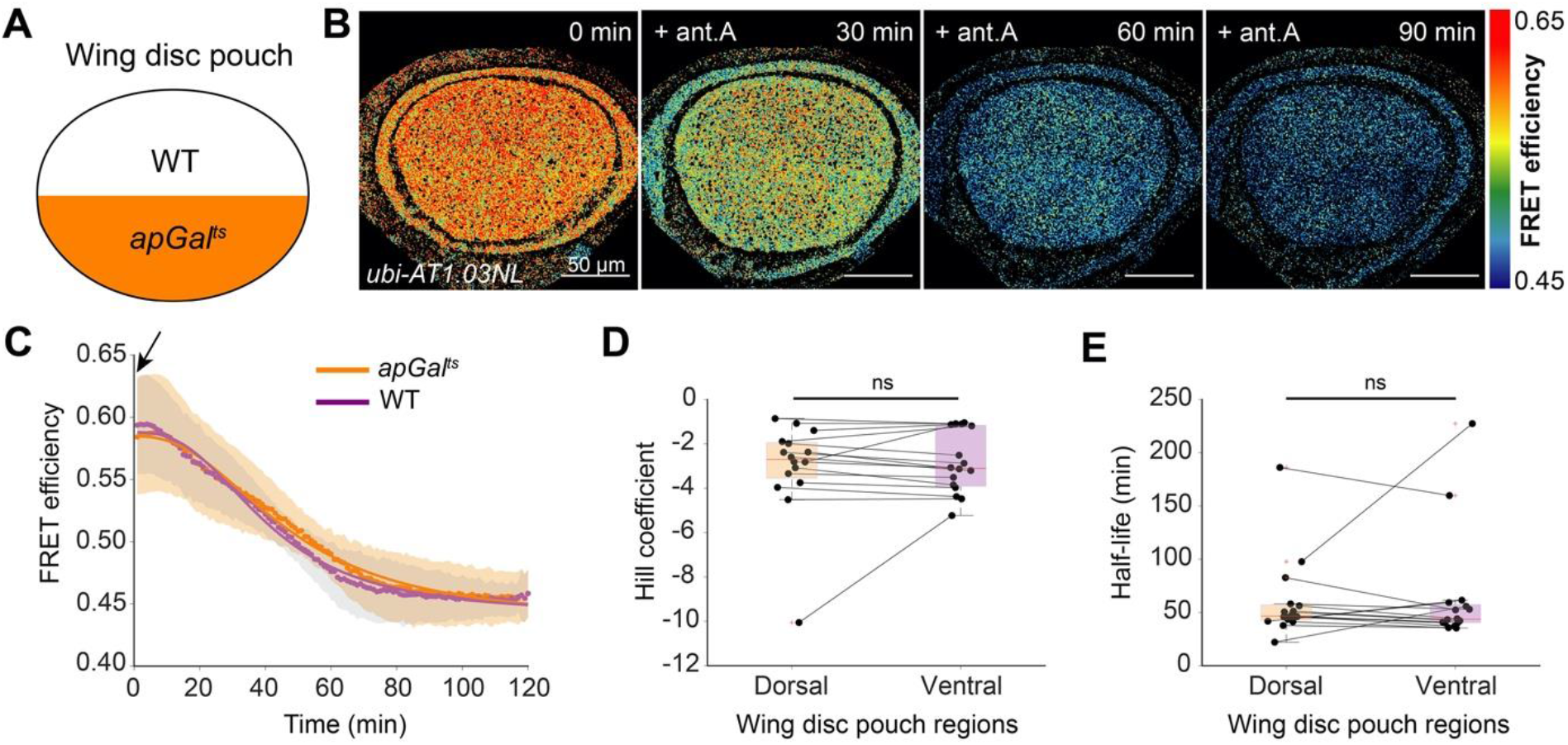
OxPhos inhibition affects ATP levels similarly in the dorsal and ventral compartments of the *apGal*^*ts*^ genetic background (without a UAS construct). (A) Schematic representation of *apGal*^*ts*^ expression in the dorsal compartment; ventral compartment serves as an internal control (WT). (B) Mean FRET efficiency measured over time in the dorsal and ventral compartments. Shaded regions indicate SD; solid lines illustrate a fit of the mean data. (C) Timelapse montage of ATP sensor FRET efficiency in the wing pouch after 10 μM antimycin A (ant.A) addition in *apGal*^*ts*^>WT wing discs. (D, E) Fit parameters of individual time traces for dorsal and ventral compartments. Black lines connect the corresponding regions of the same disc. ns = not significant, using Kruskal-Wallis test (n=16). Black arrow indicates the addition of the drugs, and small colored dots represent the averages at each timepoint.

**Extended data S4.**
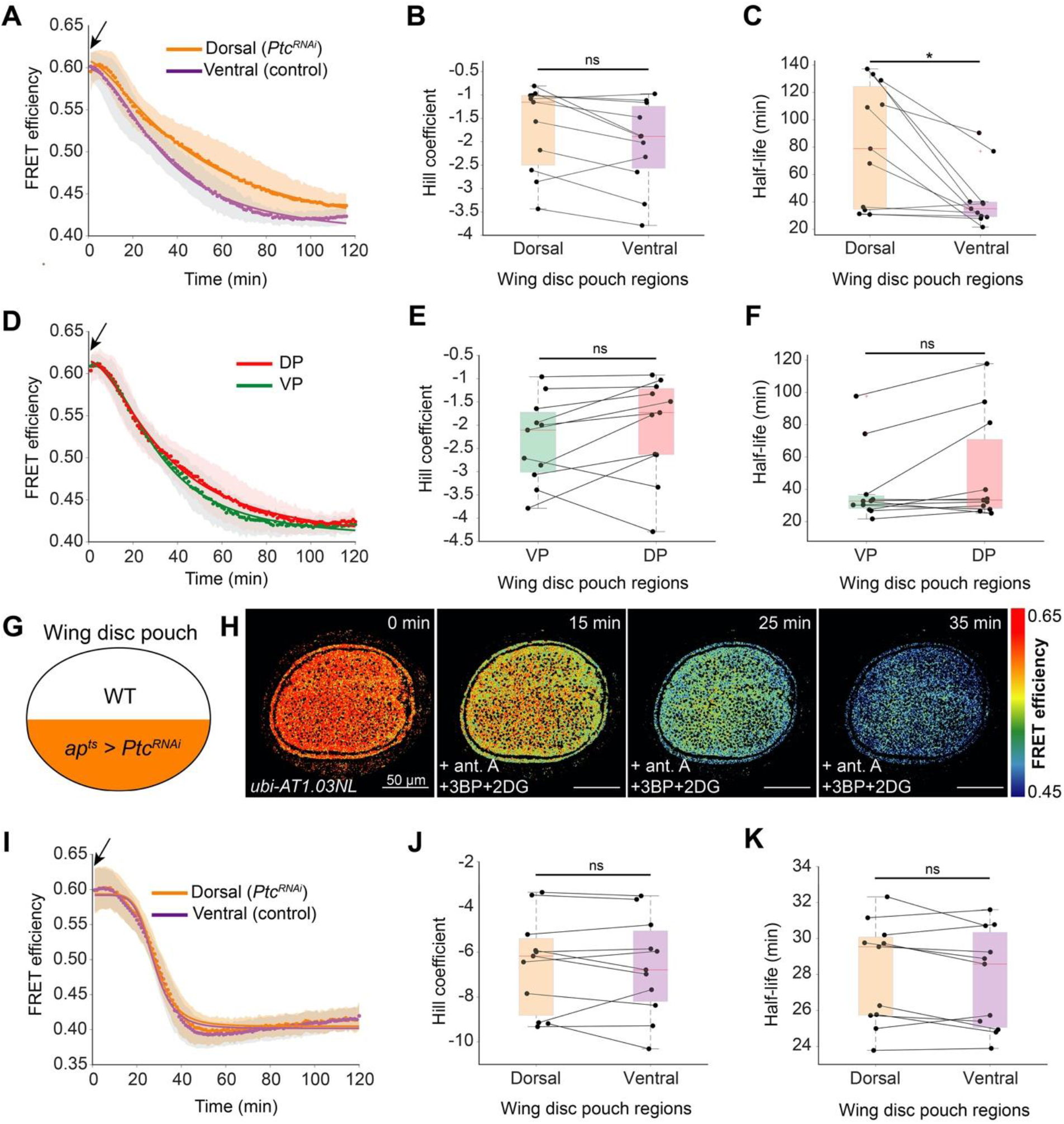
Extended regional analysis of *Ptc*^*RNAi*^ upon OxPhos inhibition alone or combined with glycolysis inhibition. (A, D) Mean FRET efficiency measured over time in the dorsal and ventral compartments (A) or the ventral posterior (VP) and dorsal posterior (DP) sub-compartments (D) of the wing pouch in *apGal*^*ts*^>*Ptc*^*RNAi*^ upon OxPhos inhibition. Shaded regions indicate SD; solid lines illustrate a fit of the mean data. Black arrows indicate the addition of antimycin A, and small colored dots represent the averages at each timepoint. (B-C, E-F) Fit parameters of individual time traces for the dorsal and ventral compartments (B-C) or the posterior sub-compartments (E-F). Black lines connect the corresponding regions of the same disc. * = p-value < 0.05, ns = not significant, using a Kruskal-Wallis test (n=11). (G) Schematic representation of *apGal*^*ts*^>*Ptc*^*RNAi*^ expression in the dorsal compartment. (H) Timelapse montage of ATP-FRET sensor efficiency after addition of antimycin A (ant. A), 3BP and 2DG. (I) Mean FRET efficiency measured over time in the dorsal and ventral compartments. Shaded regions indicate SD; solid lines illustrate a fit of the mean data. Black arrow indicates the addition of the drugs, and small colored dots represent the averages at each timepoint. (J, K) Fit parameters of individual time traces for dorsal and ventral compartments. Black lines connect the corresponding regions of the same disc. ns = not significant, using Kruskal-Wallis test (n=12).

**Figure S5.**
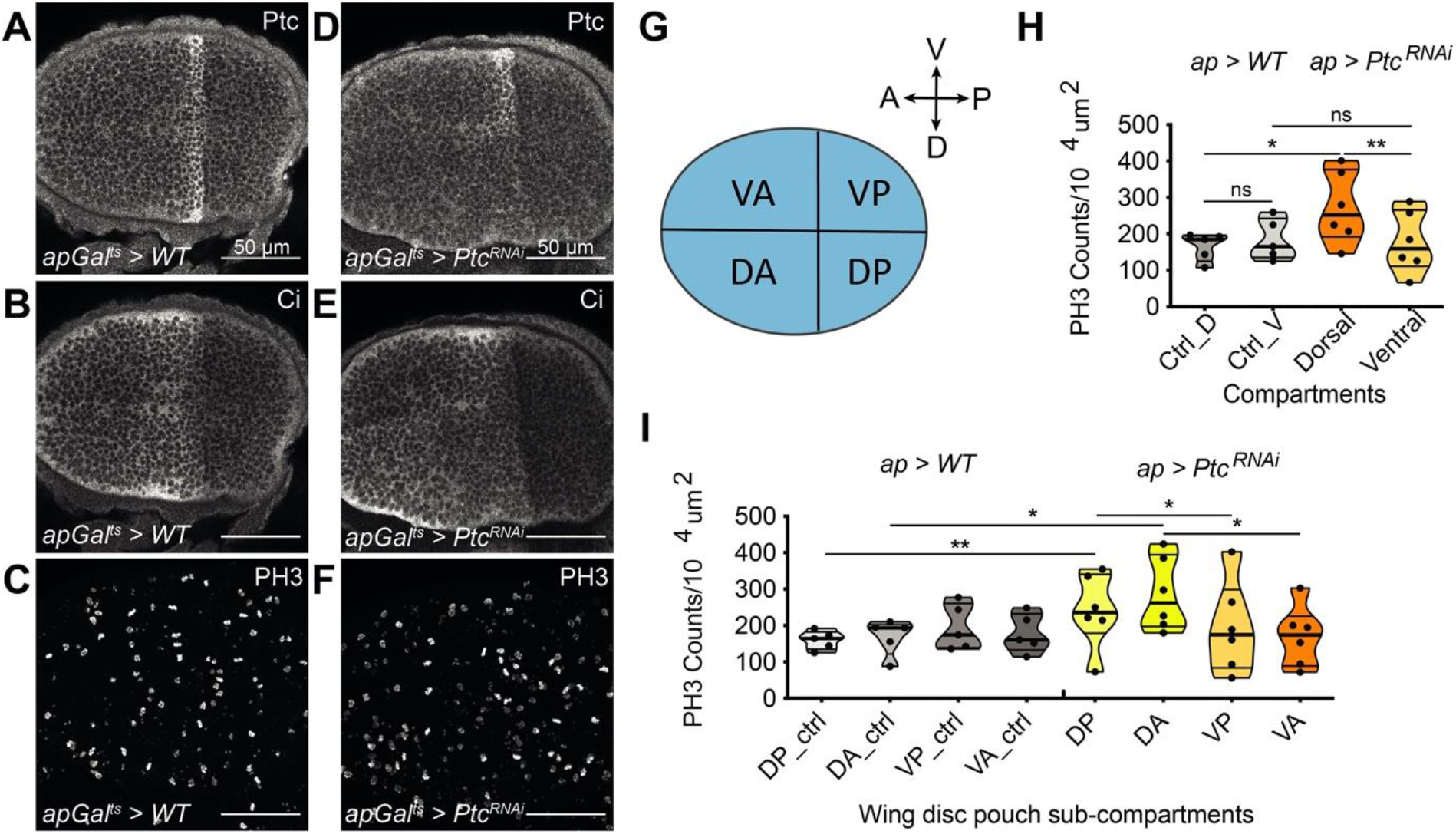
Upregulation of Hh pathway activity with *Ptc*^*RNAi*^ increases proliferation in both anterior and posterior compartments. Control (*apGal*^*ts*^>*WT*, A-C) or *apGal*^*ts*^>*Ptc*^*RNAi*^ (D-F) wing pouch stained for Ptc (A, D), Ci (B, E) or phospho-histone H3 (PH3, C, F). (G) Schematic representation of wing disc sub-compartments. V=Ventral, D=Dorsal, A=Anterior, P=Posterior. (H) Quantification of PH3-positive nuclei in the dorsal (D) and ventral (V) compartments of the pouch of control (Ctrl, *ap>WT, n=5*) and *apGal*^*ts*^>*Ptc*^*RNAi*^ (n=6) wing discs. * = p-value < 0.05, ** = p-value < 0.01 using, ns = not significant using either paired t-tests between disc compartments belonging to the same group (*apGal*^*ts*^>*Ptc*^*RNAi*^ or control) or Mann-Whitney tests for different groups (Ctrl_D vs Dorsal, Ctrl_V vs Ventral). (I) Quantification of mitotic density in different compartments of the pouch in control (Ctrl, *ap>WT*) and *apGal*^*ts*^>*Ptc*^*RNAi*^ wing discs. * = p-value < 0.05, ** = p-value < 0.01 using, ns = not significant using one-way ANOVA tests with Bonferroni post-hoc correction.

## Notes

### Competing Interest Statement

The authors have declared no competing interest.

